# Combinations of Insecticide-Treated Nets and Indoor Residual Spraying for Insecticide Resistance Management: A Modelling Exploration

**DOI:** 10.1101/2024.07.05.602226

**Authors:** Neil Philip Hobbs, Ian Michael Hastings

## Abstract

Insecticides are heavily used for the control of vectors of disease. Malaria control has been reliant on insecticide treated nets (ITNs) and indoor residual spraying (IRS). There are concerns insecticide resistance will impede malaria control. The use of insecticide resistance management (IRM) strategies is recommended. One proposed IRM strategy is the combination of ITNs and IRS. Using a mathematical model of polygenic insecticide resistance evolution, this combination strategy is evaluated. First, combinations are evaluated against ITNs alone to determine if and when combinations may be beneficial in slowing resistance evolution to the pyrethroid on the ITN. Second combinations where multiple IRS insecticides are available are compared against full-dose mixture ITNs. Results of the simulations indicate the addition of IRS to ITNs may be beneficial, providing coverage of both interventions is high. The greater number IRS insecticides available for rotation the better, however even when combinations rotate three different IRS insecticides this is still a less potent IRM strategy than deploying full-dose mixtures. In conclusion the combination of ITNs and IRS appears to offer limited benefit over full-dose mixture ITNs for an IRM perspective

## Introduction

Malaria control is reliant on insecticides for the control of transmission, with insecticide-treated nets (ITNs) and indoor residual spraying (IRS) being responsible for controlling most transmission (Bhatt et al., 2015). There is concern that increasing levels of resistance to insecticides will be problematic for transmission control (Hemingway et al., 2016). The use of insecticide resistance management (IRM) is recommended (WHO, 2012) to slow the spread of resistance both within and between mosquito populations. Modelling studies have highlighted the potential of insecticide mixtures, that is two insecticides in a single full-dose formulation, as being effective for the slowing the spread of resistance (N. Hobbs et al., 2023; Levick et al., 2017; Madgwick & Kanitz, 2022).

Mixture products are more complicated to develop, and therefore there are questions about how to obtain “temporal mixtures” through the fine-scale spatial deployments of insecticides. This could be through different households in the same village receiving different insecticides as micro-mosaics (Jones et al., 2023), although this was not found to be more effective than monotherapy rotations.

The addition of IRS to standard (pyrethroid-only) long-lasting insecticide-treated net (ITN) deployments has frequently been evaluated in cluster-randomised control trials (RCTs) (Pinder et al., 2015; Protopopoff et al., 2018; West et al., 2014). A meta-analysis of cluster-RCTs found combining IRS and ITNs may provide increased transmission control found (Choi et al., 2019). Cluster-RCTs are typically short in duration, lasting on a couple of years, and therefore any IRM benefit of combinations may not be seen.

In the Global Plan for Insecticide Resistance Management (GPIRM) (WHO, 2012), it was questioned what role combinations of ITNs and IRS play for IRM. Despite frequently being highlighted as an IRM strategy, this has not been explicitly explored empirically or theoretically. The MalERA consortium has further highlighted the need for combinations of ITNs and IRS to be evaluated for IRM (Rabinovich et al., 2017).

There are currently several IRS insecticides currently undergoing evaluation in experimental hut trials, for example chlorfenapyr (Ngufor et al., 2020), broflanilide (Snetselaar et al., 2021) and neonicotinoids (Fuseini et al., 2019). Experimental hut trials can be used to estimate the personal protection of indoor vector control products, but do not assess the long-term IRM performance of the products. The use of IRS can extend the number of insecticides available for deployment because most insecticides available for IRS are not available for use on ITNs. An experimental hut trial found the use of a mixture-ITN containing a pyrethroid and chlorfenapyr found the comparator strategy of a pyrethroid ITN with a chlorfenapyr IRS performed equally well at controlling mosquitoes (Ngufor et al., 2017). This experimental hut trial was not able to evaluate the IRM implication of deploying the insecticides in a single mixture or as a combination.

As combinations of ITNs and IRS can lead to both insecticides being deployed in the same household this means there is a greater possibility of mosquitoes encountering the insecticides in a single feeding cycle (Figure 1). Where combinations (ITN and IRS) and mixtures differ from one another is:

**Figure 1:**
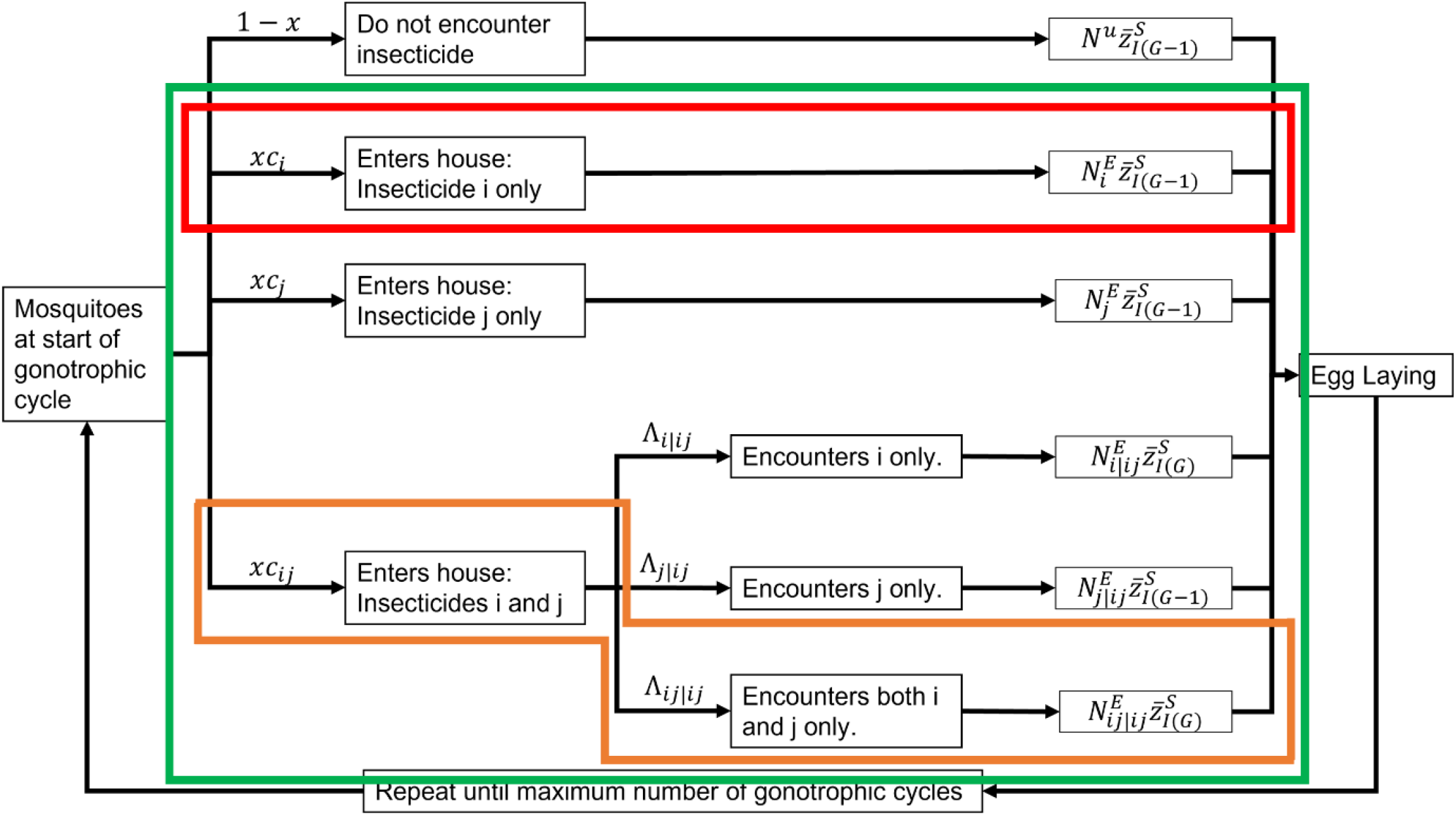
Diagram of the coverage and encounter permutations. Red box is for standard (pyrethroid-only) ITNs. Orange box is for mixtures. Green box is for combinations. Male mosquitoes only go through the selection process once.

1. Mixtures have both insecticides in the same formulation such that any mosquito contacting the mixture is expected to be simultaneously exposed to both insecticides.
2. In combinations the ITN and IRS are spatially separated even when deployed in the same house, such that a mosquito entering a house with both, may encounter just the ITN, just the IRS or both.
3. In combinations because the ITN and IRS are separate products, this means some houses may receive both, just the ITN or just the IRS.
4. As the ITN and IRS are separate products the insecticides can be switched independently of one another, which is not possible for mixtures.

The aim of this paper is to explore the role of IRS as part of an IRM strategy, and not the use of IRS as part of a transmission control strategy. This paper considers both an evaluation of adding IRS to standard pyrethroid ITN deployments, and evaluating against mixture ITNs.

## Methods

### Model Summary

Simulations are implemented using a dynamic model of insecticide selection, “polysmooth” (Hobbs & Hastings, 2024). The model assumes insecticide resistance is a polygenic trait. Selection is implemented in probabilistically, dependent on the level of resistance and efficacy of the insecticide. Female mosquitoes can complete multiple gonotrophic cycles (Figure 1), allowing them to be exposed to different insecticides throughout their lifespan. The model is written in R (R Core Team, 2020) and simulations and data analysis was performed in the R environment also. The “polysmooth” code is maintained in github repository: https://github.com/NeilHobbs/polysmooth/. When modelling the combination of IRS and ITN there is a large amount of operational and behavioural parameter space (see Equations 10a and 10c in (Hobbs & Hastings, 2024) for detail). To summarise, when deploying ITNs and IRS some houses may receive only the ITN (*c*_*i*_), some houses to only have an IRS (*c*_*j*_) and some houses to have both IRS and ITN (*c*_*ij*_).

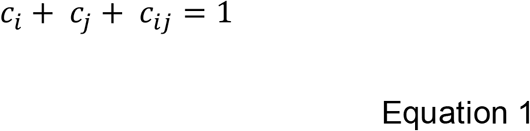

Added to this is then the mosquito behaviour variation of mosquitoes entering a house with IRS and ITN (*c*_*ij*_). Where they can encounter just the ITN 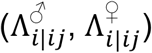, just the IRS 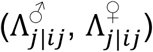 or both the ITN and IRS 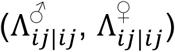:

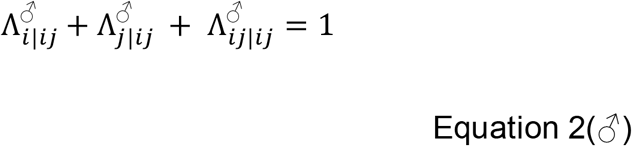

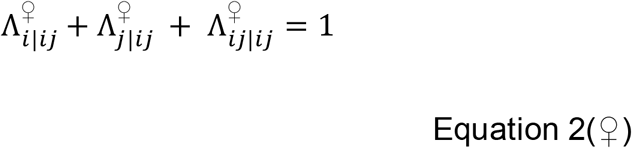

Due to this, scenarios are simplified to improve of interpretability of results. The combination strategy is evaluated in two scenarios. Scenario 1 involves adding IRS to standard ITN deployments looking at the impact of insecticide resistance. Scenario 2 looks at comparing mixture ITNs versus combinations, where the combinations are allowed an increasing number of IRS insecticides.

### Methods Scenario 1: Adding IRS to Standard (Pyrethroid-Only) ITN Deployments

The first scenario evaluates combining ITNs and IRS from a purely IRM perspective. The IRM benefit of adding IRS to ITN deployments was explored by comparing the combination (IRS + ITN) strategy against deploying just ITNs over a timeframe of 200 mosquito generations (∼20 years). In this comparison, the IRS is being added on top of ITN distributions.

A key requirement is to balance coverages between the two strategies. The total coverage of ITNs is *c*_*i*_+ *c*_*ij*_ and the total coverage of IRS is *c*_*j*_ + *c*_*ij*_. The input values of *c*_*i*_, *c*_*j*_ and *c*_*ij*_ are the same for both the combination and ITN only simulations. However, for the ITN only simulations, the efficacy of insecticide *j* (the IRS) is set to zero and is therefore absent. As the IRS insecticide is absent from the ITN only simulations, the conditional encounter values 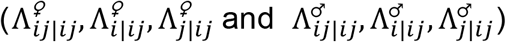, were set such that all mosquitoes entering a house with an ITN encountered the ITN 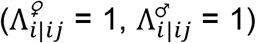.

The initial resistance to the pyrethroid insecticide on the ITN is set at 0.5, 10, 20, 50 or 80% bioassay survival. The initial resistance to the IRS insecticide is set at 0.5% bioassay survival. The two strategies are evaluated over a wide parameter space (Table 1). A total of 10,000 parameter sets were randomly generated using Latin hyperspace sampling (Carnell, 2020). The gonotrophic cycle length was fixed at 3 days, and the maximum number of gonotrophic cycles was 5. Daily survival was set between from 0.66 to 0.95 (Matthews et al., 2020).

**Table 1:**
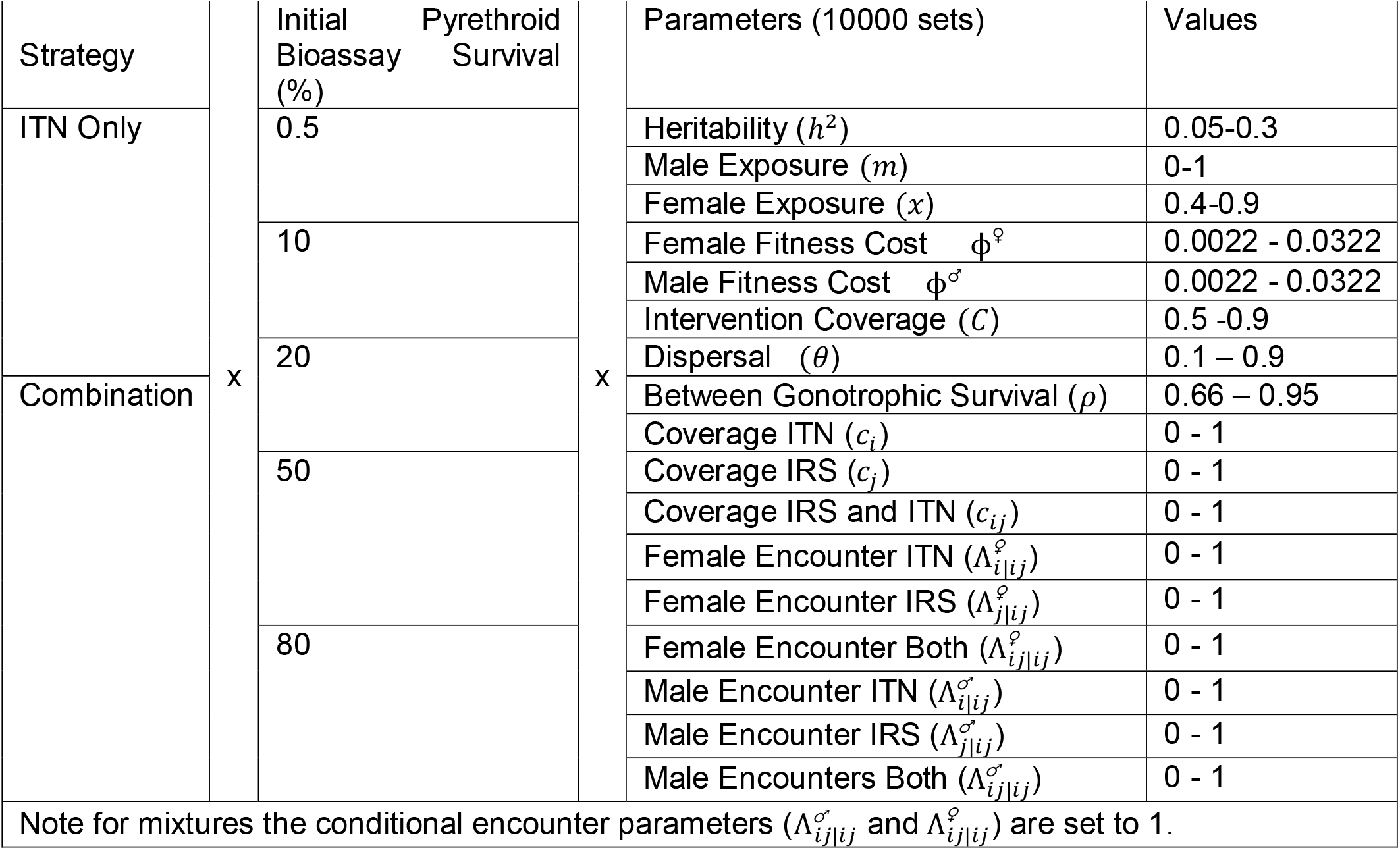
Simulation Design for Scenario 1.

The outcomes are the differences in resistance to the ITN and IRS insecticides as measured in bioassays after 200 generations (∼20 years):

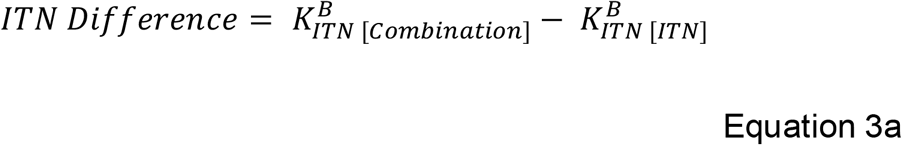

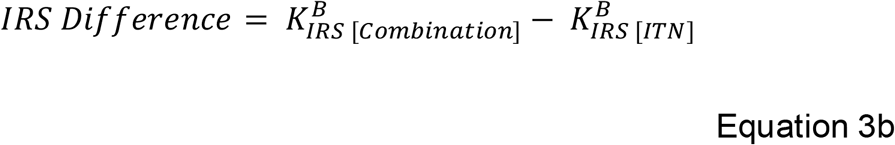

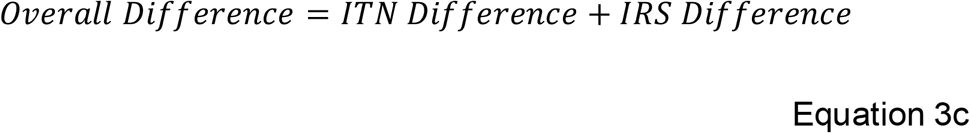

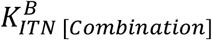and 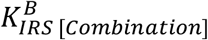 are the bioassay survivals to the ITN and IRS for the combination simulations after 200 generations. 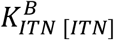 and 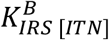 are the bioassay survivals to the ITN and IRS for the ITN only simulations after 200 generations. 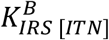 is 0.5% as the IRS is never deployed and there is no cross resistance, and this is the starting bioassay survival in the combination simulations. The primary analysis plots the distributions of *ITN Difference, IRS Differece* and *Overall Difference*. Sensitivity analysis was conducted using generalised additive models of the *ITN Difference* smoothed over each of the randomly sampled parameter inputs (Table 1).

### Methods Scenario 2: Combinations with Multiple IRS Insecticides vs Mixtures

The second scenario concerns the deployment of combinations versus the deployment of mixtures (full-dose), with the aim to determine how many IRS insecticides are required to be available to be equivalent to a full-dose mixture. Full-dose mixtures have been found to be the upper-benchmark for IRM strategy comparisons when two insecticides are available (Hobbs & Hastings, 2024a; Madgwick & Kanitz, 2022). The combinations simulations were run allowing for the inclusion of 1 (*j*), 2 (*j,k*) or 3 (*j, k, l*) IRS insecticides with rotations every 10 generations (∼yearly). For the 1 IRS simulations, insecticide *j* was deployed continuously.

To balance the coverages and exposures between the mixtures and combination simulations (i.e., that some households in the combinations simulations only receive the IRS), the proportion of mosquitoes unexposed (1-*x* for females) and (1 – *mx* for males) was balanced such that for the mixture only simulations the value of *x* was modified to account for mosquitoes which would enter IRS only houses:

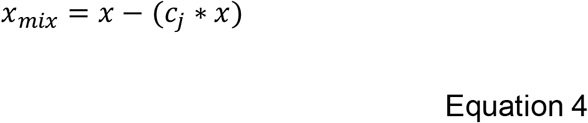

The simulations were for 200 generations (∼20 years). For each simulation all insecticides started at 0% bioassay survival, with a fixed standard deviation of the polygenic resistance score of 50. The maximum number of gonotrophic cycles was 3, (as novel insecticides are used fewer gonotrophic cycles are needed as a greater proportion of eggs are laid in the first gonotrophic cycle), with a 3 day duration and 0. 8 daily survival probability (Matthews et al., 2020). Biological parameter sets of female insecticide exposure, male insecticide exposure, intervention coverage and dispersal were created using Latin hypercube sampling (Carnell, 2020), with 500 parameter sets. Fitness costs were not included. Each of these parameter sets was then run over 60 coverage (*c*_*i*_, *c*_*j*_, *c*_*ij*_) and conditional encounter (Λ_*i*|*ij*_,Λ_*j*|*ij*_, Λ_*ij*|*ij*_) parameter sets.

The outcomes evaluated were:

- The end resistance to the ITN insecticide (*i*) insecticide after 200 generations (∼20 years). For all simulations insecticide *i*is always deployed, and therefore this gives an indication of how well this insecticide was protected:

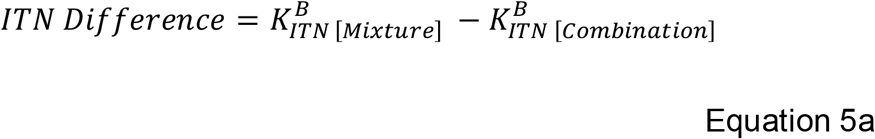
- The mean resistance to the partner insecticide (all insecticides that are not the ITN insecticide, *i*).

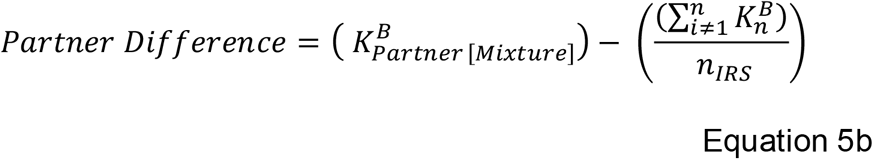

where *n*_*IRS*_ is the number of IRS insecticides in the simulation.
- The difference in the total amount of resistance to all insecticides after 200 generations (∼ 20 years). This acts as a measure of the total amount of resistance:

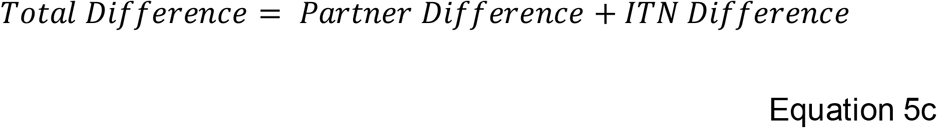

## Results

### Results Scenario 1

Figure 3 shows the difference in the level of resistance (measured as bioassay survival) after 200 generations (∼20 years) between standard (pyrethroid-only) ITNs versus a combination of a standard (pyrethroid-only) ITN and a non-pyrethroid IRS. The results of the addition of an IRS can lead to an increase the end amount of resistance to the pyrethroid (Figure 2, top plot, blue bars) will at first seem counter-intuitive. However, this can be explained by the IRS being applied in addition to the ITN. Therefore, there are households in the combination simulations receiving only the IRS which would have been untreated in the ITN only simulations. This means that in the intervention site in the combinations strategy additional mosquitoes are killed due to more houses being treated. This therefore means that the unexposed to ITNs group is smaller and the comparative sizes of the exposed groups is higher and as such are less diluted by unexposed individuals. Therefore, while there is expected to be higher levels of population control (due to increased killing) in the combinations strategy this may counter-intuitively come at the expense of faster evolving resistance to the pyrethroid. The overall “total resistance” change for the combinations was often found to be higher than for the ITN only simulations. However, this would be due to more insecticide being deployed in the combination simulations. This highlights a clear issue in evaluating IRM strategies.

**Figure 2:**
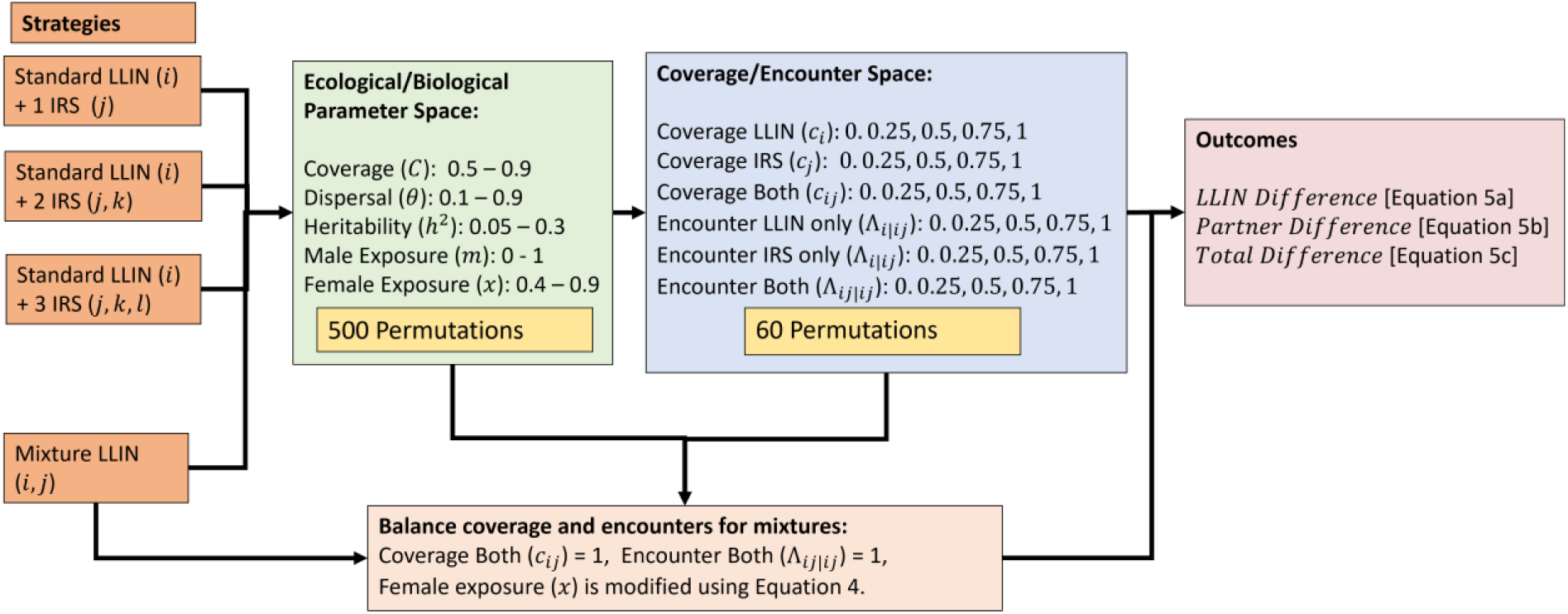
Schematic overview of Scenario 2. For each strategy evaluation, 500 randomly sampled ecological/biological parameters are used and 60 permutations of the coverage and encounter parameters, for a total of 30,000 simulations each strategy. The same inputs are used for each strategy to allow for direct comparison. To balance coverages between the Mixture ITN simulations and combinations simulations the female exposure parameter is modified using Equation 4. This also modifies male exposure as male exposure is implemented as a proportion of female exposure.

**Figure 3:**
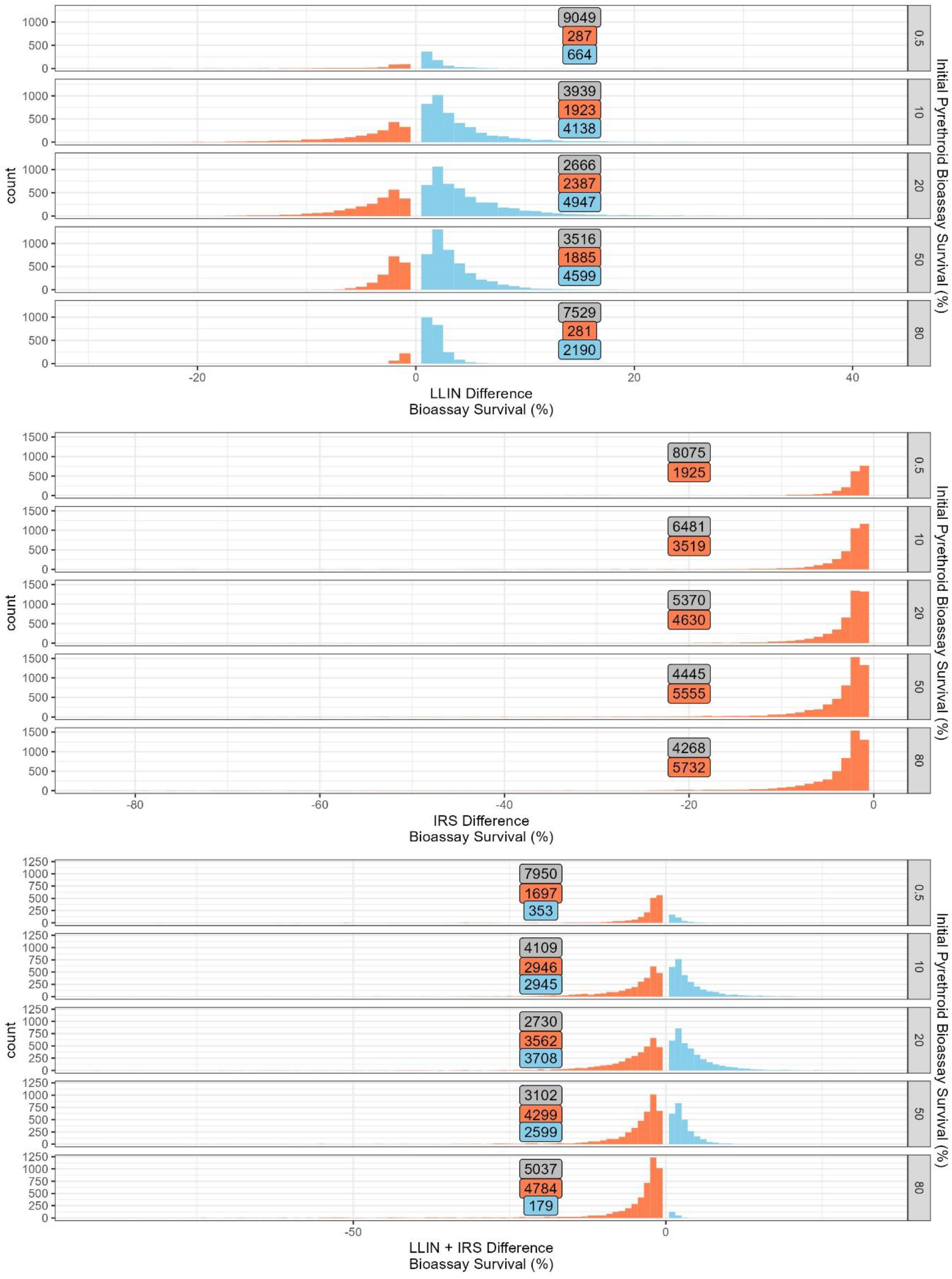
Insecticide Resistance Management impact of the addition of IRS deployments to ITN deployments. Negative values (red bars/boxes) indicate the ITN only simulations ended with lower levels of resistance. Positive values (blue bars/boxes) indicate the combination simulations ended with lower levels of resistance. Grey boxes are the number of draws. Top plot is ITN insecticide, middle plot is the IRS insecticide, and the bottom plot is for both insecticides. Each panel (top-bottom) is the initial amount of resistance measured as bioassay survival to ITN insecticide.

In general, the comparative advantage of adding IRS to ITNs is when both deployed in the same house and both insecticides are encountered (Figure 4). Under these circumstances combinations perform as a mixture. Additionally, combining ITN and IRS was found to be more especially beneficial when the “natural” daily survival was high, indicating the combination strategy was benefiting from a temporal mixture whereby mosquitoes encounter different insecticides in different gonotrophic cycles.

**Figure 4:**
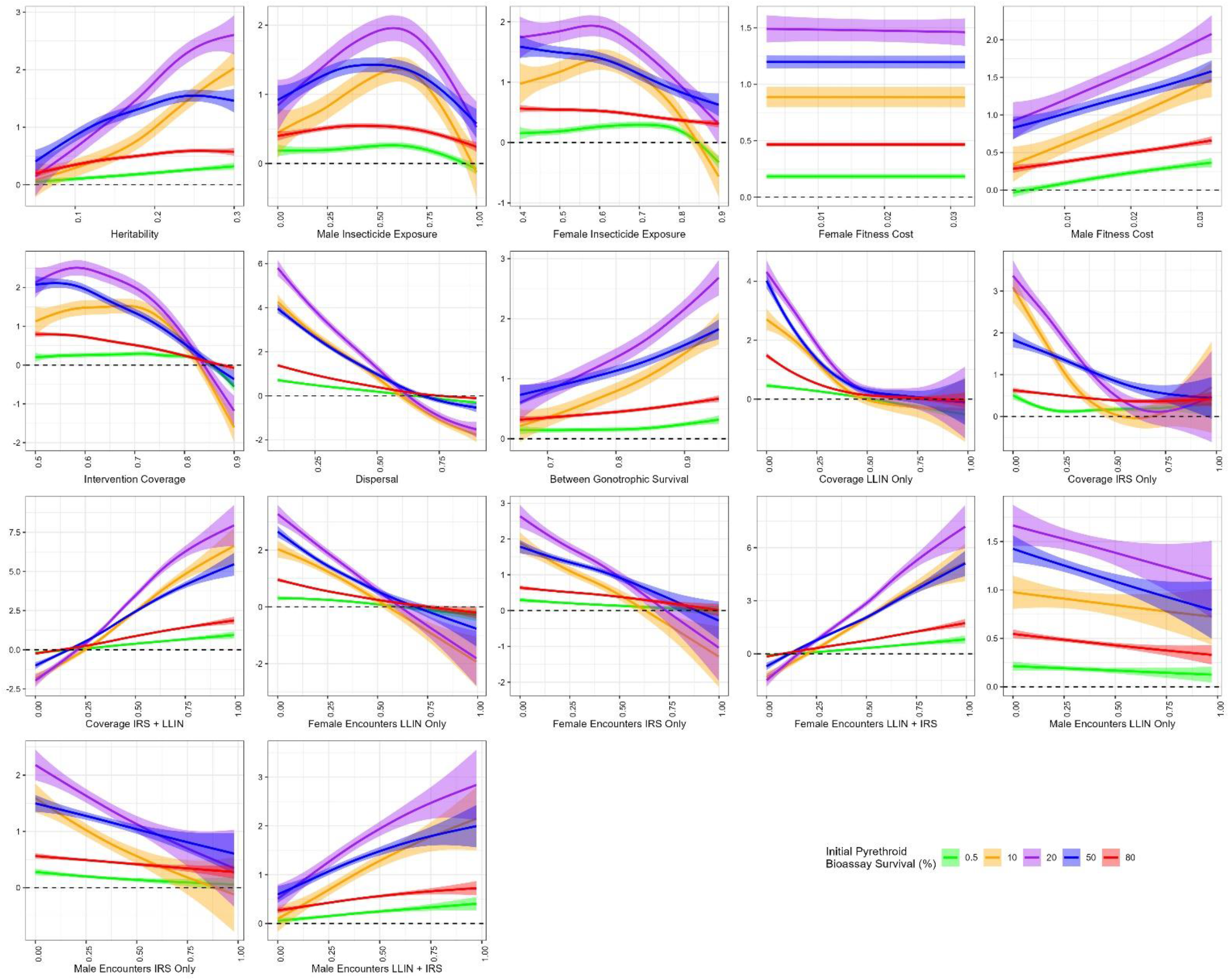
Sensitivity analysis using generalised additive models of the difference in the final bioassay survival to in the ITN insecticide smoothed over each parameter. Positive values (above the dashed line) indicate the combinations strategy performed better. Negative values (below the dashed line) indicated the ITN only strategy performed better.

### Results Scenario 2

Scenario 2 looked at the advantage of being able to rotate different IRS insecticides when compared against mixture-ITNs. Even with 3 IRS insecticides available, the use of full-dose mixture ITNs still performed better as an IRM strategy. The benefit of adding more IRS insecticides is due to these partner insecticides having less selection (due to reduced time in deployment), meaning they are maintained at lower levels of resistance (Figure 5, middle column plots). However, for the insecticide which remained deployed throughout (insecticide *i*, the ITN insecticide), less protection was supplied to this insecticide than if deployed as a mixture ITN (Figure 5, left column plots). Sensitivity analysis highlights the use of multiple IRS insecticides in combination is best when coverage of both the IRS and ITN is high and that both the insecticides are encountered in a single gonotrophic cycle (Figure 6) and that adding increasing the number of IRS insecticides is beneficial.

**Figure 5:**
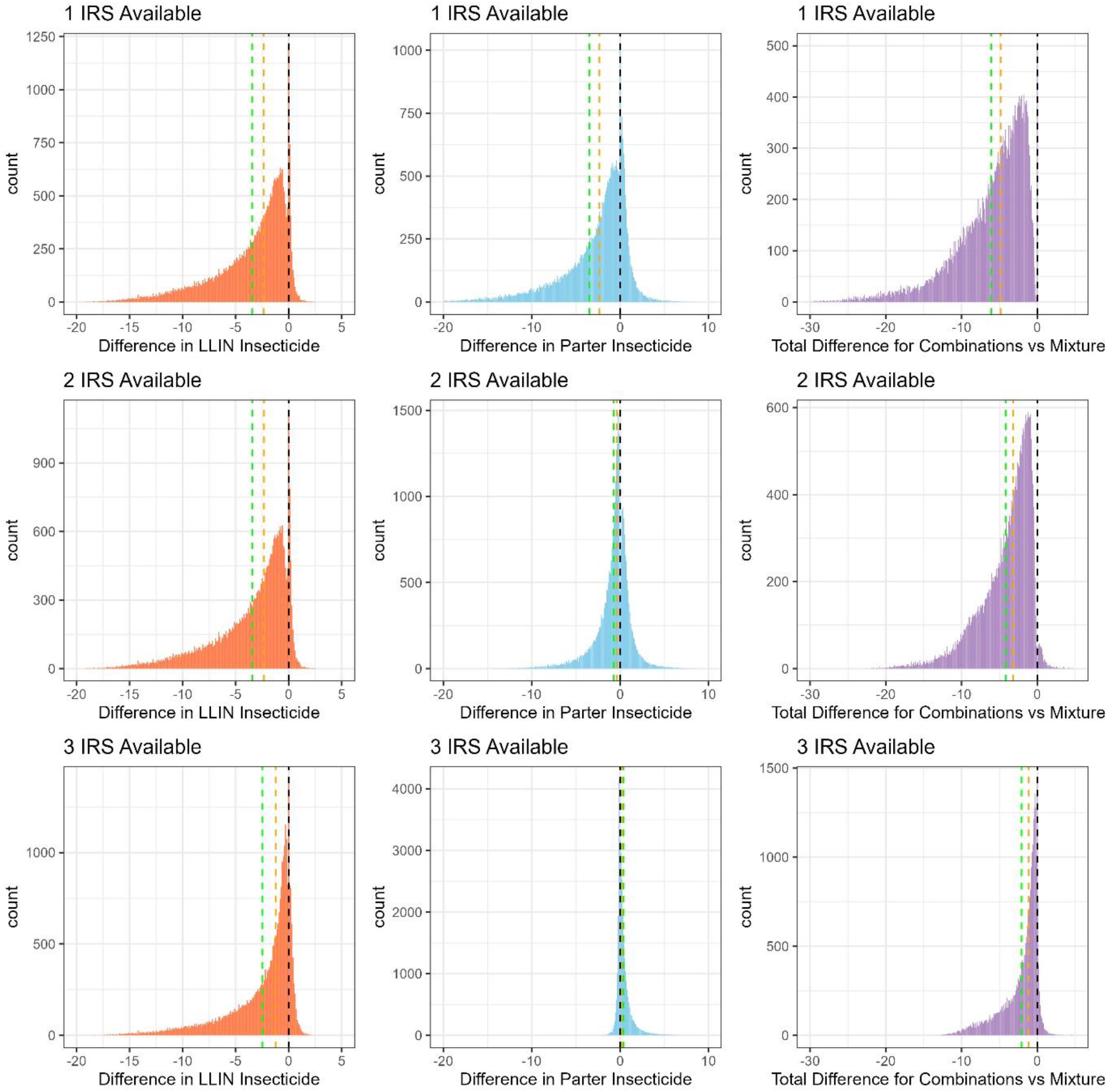
Histogram of the impact of the number of IRS insecticides available on IRM performance compared against mixtures. Top row: Only 1 IRS insecticide is used. Middle row: 2 IRS insecticides are used. Bottom row: 3 IRS insecticides are used. Left plots (light red) are for the difference in the ITN insecticide (*i*). Middle plots (light blue) difference in the partner insecticides. Right plots (purple) is the total difference. Negative values indicate the mixture performed better; positive values indicate the combination performed better. Green dashed line = mean. Orange dashed line = median.

**Figure 6:**
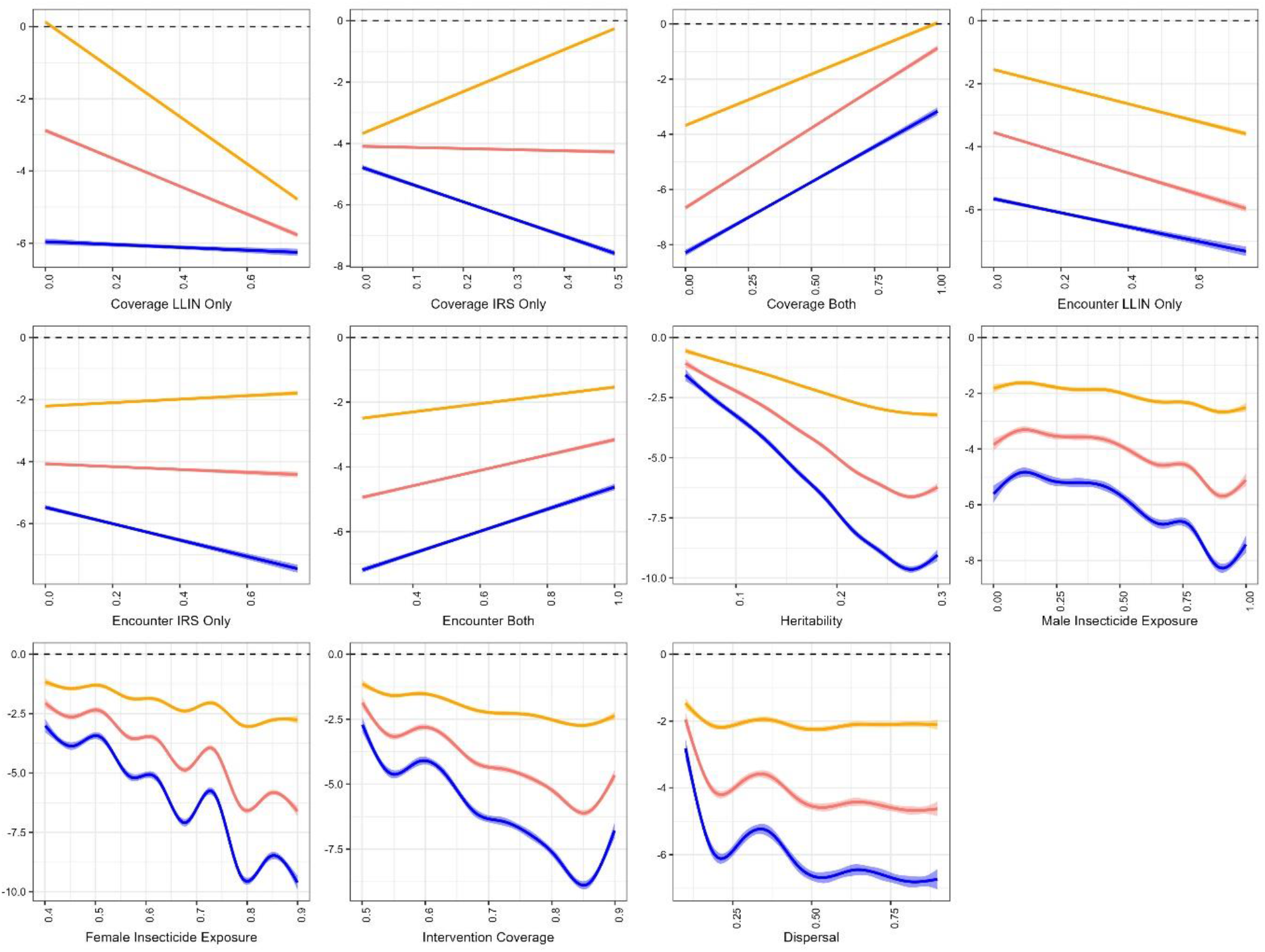
Sensitivity analysis using linear and generalised additive models for the parameter inputs on the *Total difference*. The y-axis is *Total Difference* calculated from Equation 5c. Linear models were used for Coverage/Encounter ITN/IRS/Both parameters. And generalised additive models were used for the Heritability, Exposure, Intervention Coverage, Dispersal parameters. Positive values (above the dashed line) indicate the combinations strategy performed better than the comparator Mixture ITN. Negative values (below the dashed line) indicated the ITN only strategy performed better. Blue is 1 IRS, red is 2 IRS and orange is 3 IRS insecticides.

## Discussion

The use of combinations (ITNs and IRS) as a part of IRM strategies has been highlighted as an issue requiring urgent evaluation (Rabinovich et al., 2017; WHO, 2012). Using a mathematical model which assumes a polygenic basis of resistance, “polysmooth” (Hobbs & Hastings, 2024) two key areas of this evaluation have been conducted using computational simulations. The first looking at the ability for combinations to protect standard ITNs as part of an IRM. The second, looking at the IRM value of mixture ITNs versus the use of combinations when multiple different IRS insecticides are available.

In general, it was found that adding IRS on-top of standard ITNs was generally better than not deploying the IRS (Figure 3). However, there are circumstances where adding IRS was notably detrimental, involving high levels of resistance to the standard ITN and low coverage ITN with high coverage IRS. The finding that adding IRS to ITN campaigns can increase the rate at which resistance evolves to the ITN insecticide is counterintuitive but explainable by understanding that the IRS is added on top of ITN distributions, effectively increasing coverage (and reducing the number of unexposed parents). A similar impact of adding larvicide deployments to ITN deployments may well be expected. However, with both of these, while there may be worse IRM outcomes, it would be expected there is better disease control as a result of killing more mosquitoes.

Deploying IRS has some clear benefits from an IRM perspective, some of which were not explicitly examined in the presented model. First, is the use of combinations can increase in the number of different insecticides and insecticide classes available, although the number of IRS formulations which need to be available should be high. Second, the shorter deployment durations of IRS can lead to faster and more frequent rotations. This can mean that failing insecticides can be withdrawn sooner, before there is an impact on disease control. Thirdly, IRS insecticides are deployed separately from ITNs, meaning these insecticides can then be switched independently of the insecticides used on ITNs. Forth is that IRS generally used to target short, yet intense, transmission seasons. IRS insecticides are therefore absent for the periods of lower transmission, which will mean a reduction in the insecticide selection pressure during a period when doing so is less impactful on disease transmission.

One key drawback of the combinations strategy, is that IRS is an expensive intervention to implement, and non-pyrethroid IRS formulations are considerably more expensive than pyrethroid IRS formulations (Coleman et al., 2021). The use of pyrethroid IRS has reduced substantially over the past decade (Tangena et al., 2020). The finding that full-dose mixtures outperform combinations even when 3 IRS insecticides are available, supports the roll-out of mixture ITNs.

Experimental hut trials used to evaluate the combination of ITNs and IRS (e.g. (Ngufor et al., 2011; Okumu et al., 2013; Yewhalaw et al., 2022) typically only report the ability to kill or repel mosquitoes which can be combined to give a total measure of insecticide efficacy (Briët et al., 2012). When combining ITNs with IRS, the combinations which perform best for IRM are likely to be those have minimal excito-repellency such that mosquitoes encounter both insecticides (Yakob et al., 2011).

The use of multiple IRS insecticides would necessitate the monitoring of a larger number of insecticides to ensure the tracking of insecticide resistance to ensure a failing product is not deployed. However, the ability to perform insecticide susceptibility tests is often limited due insufficient resources including insectaries and the ability to perform molecular diagnostics (van den Berg et al., 2021). In practice, the rotation of IRS insecticides is not routinely conducted (Tangena et al., 2020).

A further operational question is what role combinations play when the ITN is a mixture, leading to three insecticides being deployed simultaneously. This has been considered with synergist ITNs and IRS in experimental hut trials (Syme et al., 2022; Yewhalaw et al., 2022) where an antagonistic killing effect was found. In a cluster-RCT it was found there was also an antagonistic impact of combining synergist ITNs and IRS (Protopopoff et al., 2018). Neither the experimental hut trials nor cluster-RCT considered the long term IRM implications.

## Conclusion

The results here support the roll-out of mixture ITNs over IRS as an IRM strategy. Combinations of ITN and IRS may be beneficial in areas where only standard (pyrethroid only) ITNs are used as an interim intervention until mixture ITNs can be deployed.

## Acknowledgements

We would like to acknowledge the advice of David Weetman for help grounding the model in biological and operational relevance. We would also like to acknowledge the Vector Informatics and Genomics group at the Liverpool School of Tropical Medicine for their general critiques of the model methodology.

## Author Contributions

NPH: Data Curation, Software, Formal Analysis, Investigation, Methodology, Writing – Original Draft Preparation

IMH: Conceptualisation, Investigation, Methodology, Writing – Review & Editing, Supervision

## Data Availability Statement

Model code for the running and analysis of simulations is available from the github respository: https://github.com/NeilHobbs/polysmooth and the corresponding author upon request.

